# A frameshift mutation in the murine *Prkra* gene causes dystonia and exhibits abnormal cerebellar development and reduced eIF2α phosphorylation

**DOI:** 10.1101/2024.06.04.597421

**Authors:** Samuel B. Burnett, Allison M. Culver, Tricia A. Simon, Taylor Rowson, Kenneth Frederick, Kristina Palmer, Stephen A Murray, Shannon W. Davis, Rekha C. Patel

**Author notes:** Address Correspondence to: Rekha C. Patel, Department of Biological Sciences, University of South Carolina, 700 Sumter Street, Columbia, SC 29208 Phone: -777-4002. **Financial Disclosure/Conflict of Interest:** The authors declare no financial disclosures or conflicts of interest.

## Abstract

Mutations in *Prkra* gene, which encodes PACT/RAX cause early onset primary dystonia DYT-PRKRA, a movement disorder that disrupts coordinated muscle movements. PACT/RAX activates protein kinase R (PKR, aka EIF2AK2) by a direct interaction in response to cellular stressors to mediate phosphorylation of the α subunit of the eukaryotic translation initiation factor 2 (eIF2α). Mice homozygous for a naturally arisen, recessively inherited frameshift mutation, *Prkra^lear-5J^* exhibit progressive dystonia. In the present study, we investigate the biochemical and developmental consequences of the *Prkra^lear-5J^* mutation. Our results indicate that the truncated PACT/RAX protein retains its ability to interact with PKR, however, it inhibits PKR activation. Furthermore, mice homozygous for the mutation have abnormalities in the cerebellar development as well as a severe lack of dendritic arborization of Purkinje neurons. Additionally, reduced eIF2α phosphorylation is noted in the cerebellums and Purkinje neurons of the homozygous *Prkra^lear-5J^* mice. These results indicate that PACT/RAX mediated regulation of PKR activity and eIF2α phosphorylation plays a role in cerebellar development and contributes to the dystonia phenotype resulting from this mutation.

**Summary Statement:** This study shows, for the first time, a role of reduced eIF2α phosphorylation in DYT-PRKRA and the cerebellum development in a mouse model.

## Introduction

Dystonia is a movement disorder involving sustained muscle contractions, that can lead to painful, twisting, repetitive movements and abnormal postures (Geyer and Bressman, 2006; Bragg et al., 2011). There are multiple underlying etiologies and although several dystonia-causing mutations have been identified in various genes, the underlying pathological molecular mechanisms for most dystonia types remain unknown (Thomsen et al., 2024). The advent of next-generation sequencing technologies has made it efficient to identify the genes associated with dystonia and this list is currently growing steadily (Thomsen et al., 2024). Characterizing the genetic causes for inherited forms of dystonia offers the opportunity to investigate the underlying molecular pathomechanisms. The focus of our research presented here is DYT-PRKRA (aka dystonia 16 or DYT16), which is early-onset, generalized dystonia caused by mutations in the *Prkra* gene, which encodes the protein PACT (Patel and Sen, 1998), a stress-modulated activator of the protein kinase PKR (Patel et al., 2000). The murine homolog of PACT is termed RAX (Ito et al., 1999), and since most of the work on PACT so far has been in human cells, we will refer to *Prkra* encoded murine protein as PACT/RAX.

PKR is a ubiquitously expressed double-stranded RNA (dsRNA)-activated protein kinase (Meurs et al., 1990; Garcia et al., 2007) active under cellular stress conditions such as viral infections, oxidative and endoplasmic reticulum (ER) stress, and serum or growth factor deprivation (Ito et al., 1999; Patel et al., 2000). In virally infected cells, PKR is activated by direct interactions with dsRNA, a viral replication intermediate for many viruses (Barber, 2001) but in the absence of viral infections, other stress signals activate PKR via its protein activator PACT/RAX (Patel and Sen, 1998) in a dsRNA-independent manner. Two evolutionarily conserved dsRNA binding motifs (dsRBMs 1 and 2) in PKR allow for its interactions with dsRNA (Feng et al., 1992; Green and Mathews, 1992; Patel and Sen, 1992) as well as its dsRNA-independent interaction with PACT/RAX (Peters et al., 2001; Huang et al., 2002) and other regulatory proteins (Chang and Ramos, 2005). Upon binding dsRNA or PACT/RAX, PKR’s is activated via a conformational change and autophosphorylation (Nanduri et al., 1998; Cole, 2007). In the absence of stress, PKR stays inactive via direct interactions with the transactivation response element (TAR) RNA binding protein (TRBP) mediated by the dsRBMs of each protein (Benkirane et al., 1997; Laraki et al., 2008). TRBP inhibits PKR via the formation of both TRBP-PACT/RAX and TRBP-PKR heterodimers (Daher et al., 2009; Singh et al., 2011; Singh and Patel, 2012). Our previous work has established that the regulation of PKR activation in response to stress depends on shifting the PKR inhibitory (PACT/RAX-TRBP and TRBP-PKR) interactions to PKR-activating (PACT/RAX-PKR and PACT-RAX/PACT-RAX) interactions in response to the stress signal (Daher et al., 2009). This is regulated by stress-induced PACT/RAX phosphorylation, which dissociates PACT/RAX from TRBP and allows for its interaction with PKR (Singh et al., 2011; Singh and Patel, 2012).

PKR is one of the four protein kinases that regulate the integrated stress response (ISR), an evolutionarily conserved pathway activated by a diverse set of stress signals in eukaryotic cells. ISR works to restore the cellular homeostasis (Pakos-Zebrucka et al., 2016) by reducing the rate of general protein synthesis while allowing a selective synthesis of proteins involved in cellular recovery. The central regulatory step in this pathway is the phosphorylation of the α subunit of eukaryotic translation initiation factor 2 (eIF2α) on serine 51 by one of the four serine/threonine kinases (Donnelly et al., 2013; Taniuchi et al., 2016). Phosphorylation of eIF2α prevents the formation of the ternary complex required for translation initiation, leading to a decrease in overall protein synthesis while allows translation of selected mRNAs that have short upstream open reading frames in their 5’UTR and encode proteins that aid in cellular recovery (Wek, 2018). While transient eIF2α phosphorylation promotes cellular survival, prolonged eIF2α phosphorylation is pro-apoptotic due to transcriptional upregulation as well as preferential translation of pro-apoptotic proteins (Donnelly et al., 2013). Thus, although ISR is primarily a response to restore cellular homeostasis and promote survival, exposure to severe or prolonged chronic stress drives signaling towards cellular death.

DYT-PRKRA is an early-onset movement disorder characterized by progressive limb dystonia, laryngeal and oromandibular dystonia and parkinsonism (Camargos et al., 2008). Currently, ten PACT mutations leading to DYT-PRKRA are reported (OMIM: DYT16 (612067) in a worldwide occurrence (Camargos et al., 2008; Seibler et al., 2008; Camargos et al., 2012; Lemmon et al., 2013; Zech et al., 2014; de Carvalho Aguiar et al., 2015; Quadri et al., 2016; Dos Santos et al., 2018; Masnada et al., 2021; Bhowmick et al., 2022). Using DYT-PRKRA patient derived lymphoblasts and other *in vitro* biochemical approaches, our lab has established that PACT mutations increase cellular susceptibility to ER stress due to dysregulated eIF2α stress response signaling (Vaughn et al., 2015; Burnett et al., 2019; Burnett et al., 2020). In agreement with our findings, the dysregulation of eIF2α signaling was also reported in DTY-TOR1A, DYT-THAP1, DYT-SGCE, and sporadic cervical dystonia (Rittiner et al., 2016; Beauvais et al., 2018; Zakirova et al., 2018). Collectively, these findings indicate a potential common pathological link among some forms of inherited dystonia.

Palmer *et al*. described a spontaneous frameshift mutation (*lear-5J*) in the murine *Prkra* gene that codes for RAX, the mouse homolog of PACT (Ito et al., 1999; Palmer et al., 2016a). This frameshift mutation results in a truncation of the PACT/RAX protein within the dsRBM2 domain (Huang et al., 2002). PACT and RAX are highly homologous differing only in 6 amino acids 4 of which are conservative changes (Patel and Sen, 1998; Ito et al., 1999). The *Prkra^lear-5J^*mice present with craniofacial developmental abnormalities, small ear size (giving the name "lear" for "little ears") drastically reduced body size, kinked tails, and ascending dystonia that progresses until becoming fatal at 3-6 weeks of age (Palmer et al., 2016a). In the present study, we undertook an initial characterization of both the *in vitro* and *in vivo* consequences of the *Prkra^lear-5J^* mutation. Our findings demonstrate that the truncated protein is present in the mouse brain but not in murine embryonic fibroblasts (MEFs), the mutant PACT/RAX protein retains its ability to interact with PKR and inhibits PKR activation. Our *in vivo* data evaluating the brains of *Prkra^lear-5J^* mice demonstrates defective foliation of the cerebellum and a dramatic reduction in dendritic arborization of Purkinje neurons. Finally, we also observed a reduction in phosphorylated eIF2α, and elevated expression of CreP, an eIF2α-specific phosphatase. Our results demonstrate that dysregulating PACT/RAX mediated eIF2α phosphorylation in mouse has severe consequences on cerebellar development and may contribute to the etiology of the observed dystonia phenotype.

## Results

### A single nucleotide insertion truncates PACT/RAX in dsRBM2 in *Prkra^lear-5J^*mice

Palmer et al. identified a recessively inherited spontaneous mutation on the BTBR T^+^ Itpr3^tf^/J genetic background, that resulted in a small body size, small ears, kinked tails, progressive dystonia or paralysis, hearing loss, and mortality (Palmer et al., 2016a). They confirmed by sequence analysis that these phenotypes arose from a single adenine insertion in codon 178 of *Prkra* gene. The frameshift caused by the insertion leads to a premature stop codon after seven extraneous amino acids within the dsRBM2, one of PACT/RAX’s functional motifs (Figure 1 A and B). The resultant protein contains the entire dsRBM1 and a partial dsRBM2 eliminating some of the residues within this domain known to be important for dsRNA-binding and protein-protein interactions. However, the truncated protein retains the entire dsRBM1, which is most crucial for dsRNA-binding and protein-protein interactions based on the previous work from our group and other labs (Peters et al., 2001; Huang et al., 2002; Chukwurah et al., 2018).

**Figure 1:**
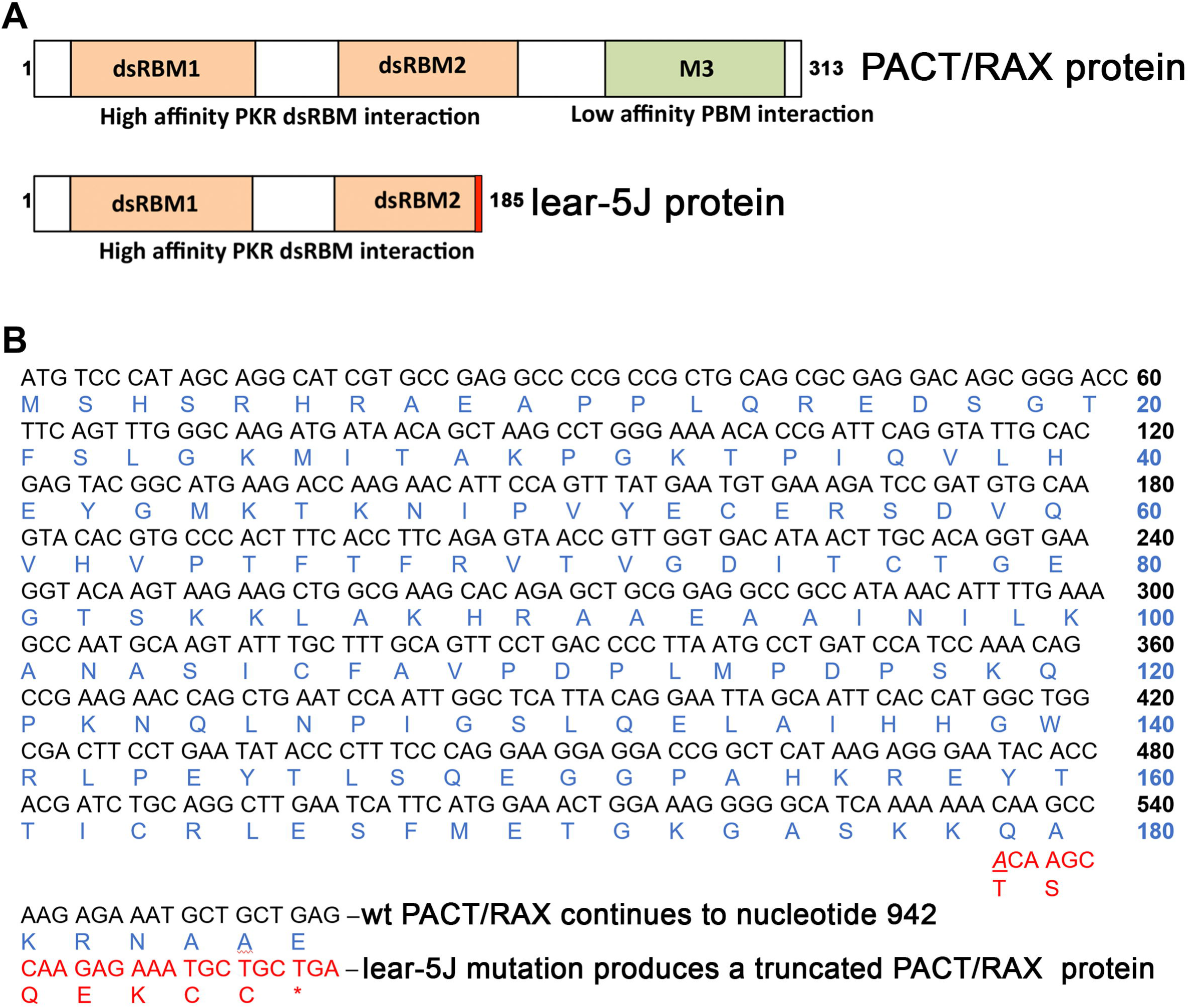
Schematic representations of the lear-5J frameshift mutation in the *Prkra* gene. (A) Functional domains of PACT/RAX and lear-5J truncated protein. Orange boxes: conserved dsRBM1 and dsRBM2 that facilitate high affinity dsRNA as well as protein-protein interactions. Green box: dsRBM3 that does not bind dsRNA but has weak binding affinity to the PACT/RAX binding motif (PBM) within the catalytic (kinase) domain (KD) of PKR. The frameshift mutation from a single nucleotide insertion results in the addition of 7 novel amino acid represented in red before the stop codon. (B) Frameshift mutation in *Prkra* ORF. Single adenine insertion at nucleotide position 534 (underlined) results in a truncated protein with the original 178 amino acids (AA) of PACT/RAX followed by 7 novel AA before a premature stop codon truncating the protein. This truncation occurs within the dsRBM2 functional domain.

### Effect of the *Prkra^lear-5J^* mutation on PACT/RAX’s dsRNA-binding and PKR interaction

To characterize how the lear-5J mutation affects the properties of the protein, we tested if the mutation affects the ability of PACT/RAX to bind dsRNA. An *in vitro* dsRNA binding assay previously well-established for PKR and PACT (Patel and Sen, 1992; Patel et al., 1996; Patel and Sen, 1998; Huang et al., 2002; Chukwurah et al., 2018) was performed using dsRNA immobilized on agarose beads and in vitro translated ^35^S methionine-labeled proteins. As seen in Figure 2 A and B, the *Prkra^lear-5J^* mutant protein shows reduced dsRNA binding in comparison to the wt PACT/RAX (Figure 2 A, lanes 2 and 4 and Figure 2 B). About 40% of the wt PACT/RAX protein bound to the dsRNA-agarose beads but only about 23% of *Prkra^lear-5J^* protein showed binding. To ascertain the specificity of the dsRNA-binding assay, we used *in vitro* translated firefly luciferase as a negative control, which does not bind to dsRNA (lanes 7 and 8). Additionally, we demonstrate the specificity of the interaction for dsRNA by adding excess dsRNA or ssRNA as competitors. As seen in lanes 5 and 6, the binding to dsRNA immobilized on beads can be effectively competed by exogenously added dsRNA but not single-stranded (ss) RNA. This ascertains the specificity of the dsRNA-binding assay, and it can be concluded that the *Prkra^lear-5J^* mutant protein shows binding to dsRNA at a reduced efficiency compared to the wt PACT/RAX protein.

**Figure 2:**
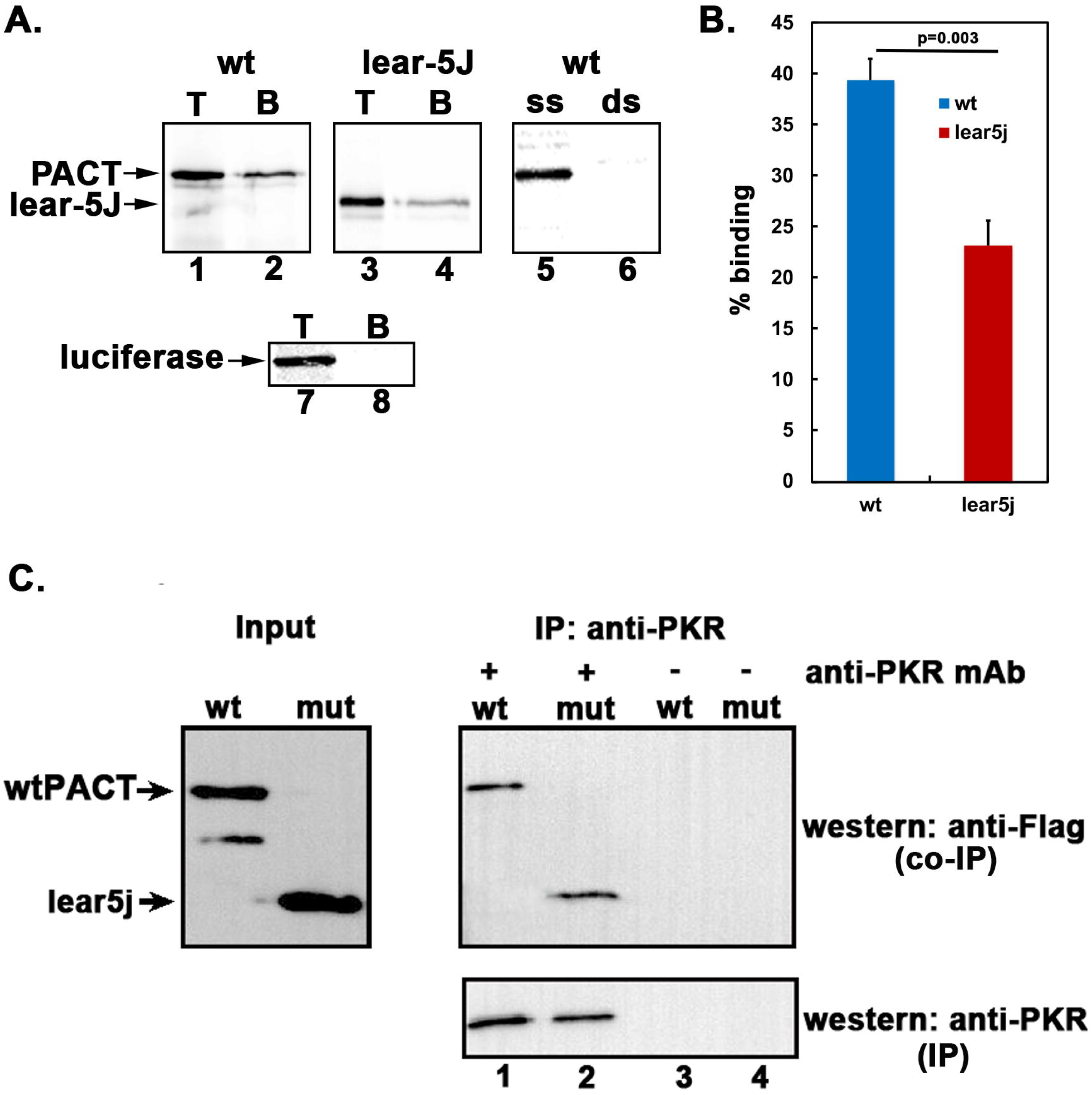
The lear-5J protein binds dsRNA less efficiently but interacts with PKR similar to wt PACT/RAX. (A) dsRNA binding assay. dsRNA binding activity of WT PACT/RAX/RAX and lear-5J truncated protein was measured by a poly(I):poly(C)-agarose binding assay with *in vitro* translated ^35^S-labeled proteins. T, total input; B, proteins bound to poly(I):poly(C)-agarose. Competition lanes 5 and 6: competition with 100-fold molar excess of single-stranded RNA (ss) or dsRNA (ds). The faint band below the parent PACT/RAX band represents products of *in vitro* translation from an internal methionine codon in lane 1. (B) Quantification of the dsRNA binding assay. Bands were quantified by phosphorimaging analysis, and % bound was calculated. Error bars: S.D. from three independent experiments. The p value (0.003) calculated using statistical analyses indicated significant difference between % dsRNA-binding of WT (blue bar) and lear-5J mutant (red bar) as indicated by the bracket. (C).Co-Immunoprecipitation of endogenous PKR and Flag-PACT/RAX or Flag-lear-5J overexpressed in HeLa cells. HeLa cells were transfected with Flag wt PACT/RAX or Flag-lear-5J in pCDNA3.1-expression constructs at 40% confluency and harvested 24-hour post transfection. Whole cell extracts were immunoprecipitated at 4°C overnight using 100 ng of anti-PKR antibody per IP. Samples were then analyzed via SDS-PAGE gel electrophoresis and western blot analysis probing for Flag tagged wt PACT/RAX or lear-5J (co-IP panel) using monoclonal anti-Flag-M2 (Sigma) antibody. To ascertain that an equal amount of protein was immunoprecipitated, blots were re-probed using an anti-PKR antibody (IP panel). Input blots without immunoprecipitation demonstrate equal amounts of each protein were present prior to IP.

Previously, our lab reported that mutating specific hydrophobic residues within PACT’s dsRBM1 significantly disrupts PACT-PKR interactions, while disrupting hydrophobic residues in dsRBM2 has minimal consequences for PACT-PKR interaction (Chukwurah et al., 2018). As the *Prkra^lear-5J^*mutation truncates the protein within PACT/RAX’s dsRBM2, we tested if the truncated protein retains its ability to interact with PKR (Figure 2 C and D). We performed co-immunoprecipitation (co-IP) assays from cells overexpressing either flag-tagged wt PACT/RAX or lear-5J proteins. Our results indicate that despite the truncation, the lear-5J protein interacts with PKR with equal binding affinity to that of wt PACT/RAX (lanes 1-2). In the absence of PKR antibody, we do not detect flag-wt PACT/RAX or flag-lear-5J thus indicating that there is no binding to the beads nonspecifically in the absence of PKR protein (lanes 3-4). Lanes 1-2 (lower panel) indicate that equal amount of endogenous PKR was immunoprecipitated in both samples while lanes 3-4 (lower panel) indicate that PKR is absent in absence of PKR antibody. Input blot (left) demonstrates equal expression of flag-wt PACT/RAX and flag-lear-5J prior to immunoprecipitation. Thus, the truncated lear-5J protein retains its ability to interact with PKR.

### Truncated lear-5J protein inhibits PKR’s kinase activity

We previously established that PACT/RAX activates PKR under conditions of cellular stress (Patel et al., 2000; Daher et al., 2009; Singh et al., 2009; Singh et al., 2011; Singh and Patel, 2012) but recombinant purified PACT/RAX protein can activate PKR robustly *in vitro* (Patel and Sen, 1998; Huang et al., 2002). Therefore, we next tested the effect of *Prkra^lear-5J^*frameshift mutation on PKR activation by performing PKR activity assays using either purified recombinant hexahistidine-tagged wt PACT/RAX or lear-5J proteins to activate PKR. Both hexahistidine-tagged wt PACT/RAX and lear-5J proteins were expressed in bacteria and purified using affinity chromatography on Ni-agarose. Based on our previous work (Patel and Sen, 1998; Huang et al., 2002), two different amounts of purified recombinant proteins were tested for activation of PKR immunoprecipitated from HeLa cells (Figure 3 A). Our results indicate that at lower concentration (400 pg), lear-5J does not activate PKR above background (lanes 1-2), whereas the wt PACT/RAX causes a robust activation of PKR (lane 4). Interestingly, lear-5J activates PKR very slightly at the higher concentration (4 ng) (lane 3). Since lear-5J has drastically reduced ability to activate PKR (lane 3) as compared to wt PACT/RAX (lanes 4-5), we next tested if lear-5J protein inhibits PKR activation brought about by dsRNA or PACT/RAX. As seen in Figure 3 B, lear-5J protein inhibits PKR activation brought about by both dsRNA (lanes 2-5) and by PACT/RAX (lanes 6-9). These results indicate that lear-5J protein inhibits PKR activation in a dose dependent manner.

**Figure 3.**
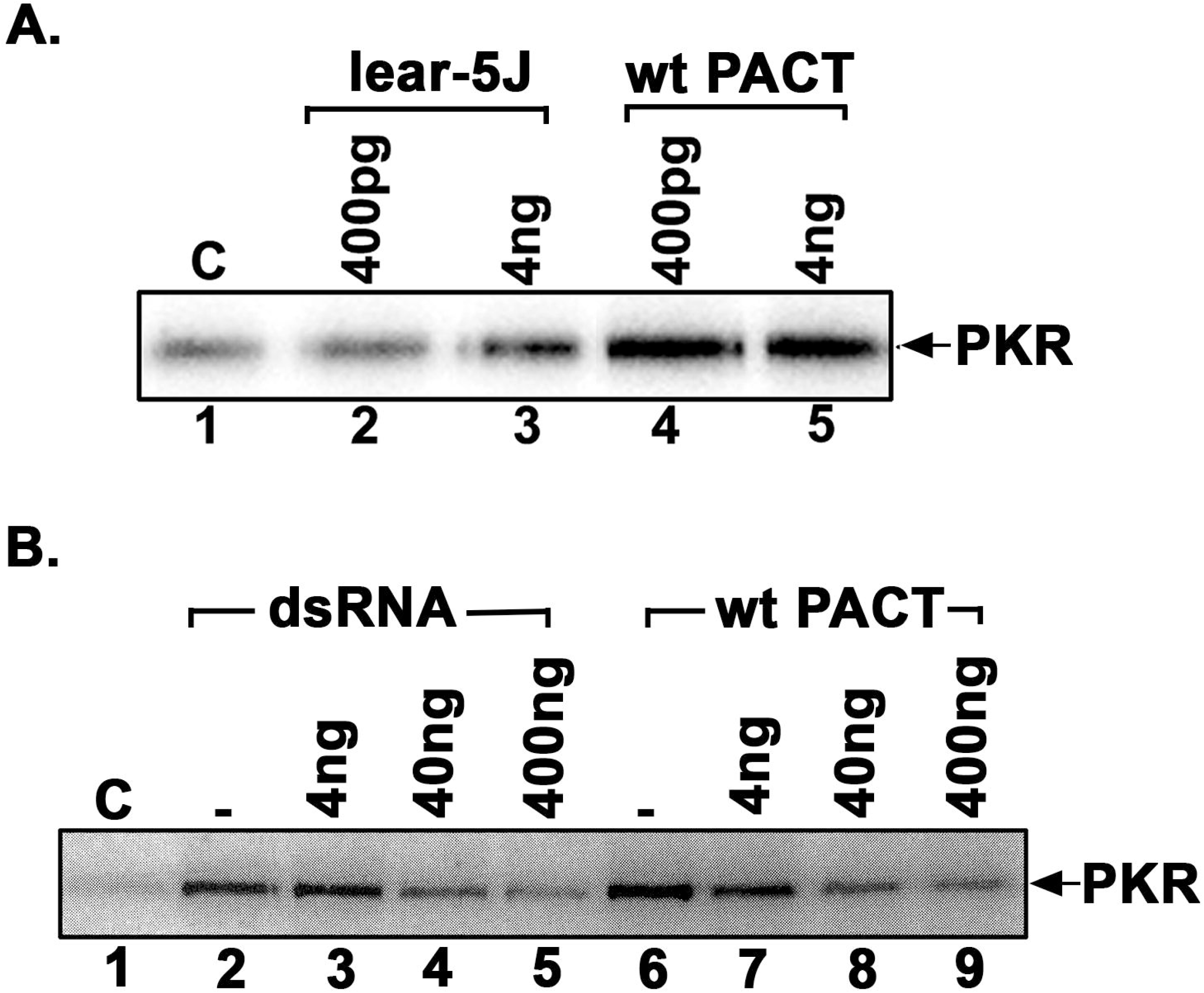
(A) Effect of lear-5J protein on PKR kinase activity. PKR kinase activity assay was performed using PKR immunoprecipitated from HeLa cells, recombinant lear-5J and wt PACT/RAX proteins, and 1 μCi of [γ-^32^P] ATP per reaction. Either pure recombinant lear-5J (left) or wt PACT/RAX (right) protein was added as activator in amounts indicated above each lane. Phosphorylated PKR was visualized after SDS-PAGE and phosphorimager analysis. (B) The truncated lear-5J protein inhibits PKR activation. PKR immunoprecipitated from HeLa cell extracts was activated with of polyI:polyC (lanes 2-5) or 4 ng of recombinant pure wt PACT/RAX protein (lanes 6-9). Increasing amounts of recombinant pure lear-5J protein (4 ng-400 ng) were added (lanes 2-9) as indicated to assess the effect on PKR activity. Lane 1: PKR activity without any added activator, lane 2: PKR activity in the presence of polyI:polyC, lanes 3-5: PKR activity in the presence of polyI:polyC and 4ng, 40 ng, or 400 ng of lear-5J protein, lane 6: PKR activity in the presence of 4ng of wt PACT/RAX protein, lanes 7-9: PKR activity in the presence of 4 ng of wt PACT/RAX and 4 ng, 40 ng or 400 ng of lear-5J protein. Phosphorylated proteins were analyzed by SDS-PAGE and phosphorimager analysis.

### Mouse embryonic fibroblasts (MEFS) isolated from *Prkra^lear-5J^*mice are resistant to ER stress-induced apoptosis

We have previously established that PACT/RAX-induced PKR activation is involved in regulating cell survival in response to endoplasmic reticulum (ER) stress (Singh et al., 2009; Vaughn et al., 2015; Burnett et al., 2020). Thus, we evaluated the response of *Prkra^lear-5J^*MEFs with that of wt (BTBR T^+^ Itpr^tf^/J) MEFs. To compare the relative apoptosis in wt and lear-5J MEFs, we used DNA fragmentation analysis in response to tunicamycin treatment. Tunicamycin (TM) treatment results in accumulation of misfolded proteins in the ER due to inhibition of protein glycosylation and triggers ER stress response. DNA fragmentation is a late marker of apoptotic cells as the DNA is cleaved by caspase-activated DNases (CADs) into nucleosomal fragments of 180 bp (Nagata et al., 2003). As seen in Figure 4 A, the wt (BTBR T^+^ Itpr^tf^/J) MEFs showed significantly high levels of DNA fragmentation in response to tunicamycin (lanes 5-7). In comparison, the *Prkra^lear-5J^* MEFs have no detectable DNA fragmentation after exposure to tunicamycin (lanes 2-4). These results indicate that *Prkra^lear-5J^* MEFs are significantly protected from ER stress-induced apoptosis.

**Figure 4.**
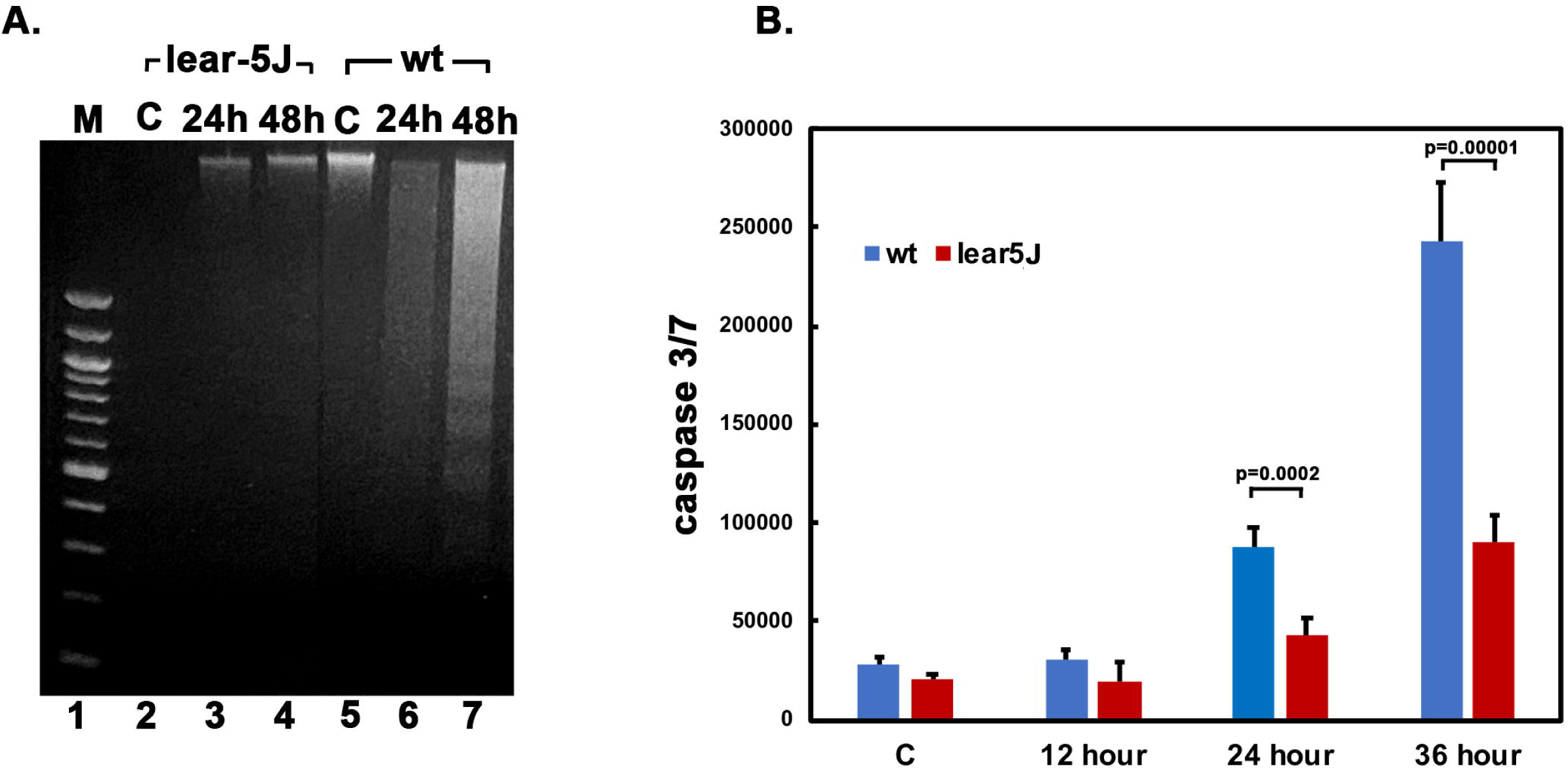
Tunicamycin-induced apoptosis is reduced in *Prkra^lear-5J^* MEFs. A, DNA fragmentation analysis. MEFs established from wt (BTBR T^+^ Itpr^tf^/J) and *Prkra^lear-5J^* mice were treated with 0.5 µg/ml tunicamycin for indicated times. Lane 1: 100 bp marker ladder, lanes 2 -4: WT (BTBR T^+^ Itpr^tf^/J) MEFs; lanes 5 -7: *Prkra^lear-5J^* MEFs.L Lanes 2 and 5: untreated cells; lanes 3-4 and 6-7: tunicamycin-treated cells. (B) Caspase-Glo 3/7 assay. MEFs established from WT (BTBR T^+^ Itpr^tf^/J) mice and *Prkra^lear-5J^* mice were treated with 0.5Lµg/ml tunicamycin for the indicated time points and caspase 3/7 activities were measured. Blue bars, WT (BTBR T^+^ Itpr^tf^/J) MEFs; red bars, *Prkra^lear-5J^*MEFs. The p values that were significant are as indicated.

To further validate these results, we performed caspase 3/7 activity assays under the same treatment conditions to measure apoptosis. In wt MEFs we detect caspase activity above control levels at 24 h which increases at 36 h post-treatment (Figure 4 B, blue bars). In contrast, the *Prkra^lear-5J^* MEFs demonstrate significantly reduced caspase activity at any of the time points post-treatment (Figure 4 B, red bars). This substantiates that the *Prkra^lear-5J^* MEFs are significantly resistant to ER stress and exhibit less apoptosis as compared to wt MEFs.

### Lear-5J protein is detectable in mouse brain but not in MEFs

The mRNAs with frameshift mutations containing an early stop codon are often degraded via nonsense mediated decay (NMD) (Carrard and Lejeune, 2023; Patro et al., 2023; Petrić Howe and Patani, 2023). We next wanted to address whether the lear-5J mutant protein and mRNA is detectable in mice, or if the expression of lear-5J is silenced partially or fully via NMD. To address this question, we isolated protein and total RNA from the brains of wt (BTBR T^+^ Itpr^tf^/J), heterozygous *Prkra^lear-5J^*, and homozygous *Prkra^lear-5J^*mice. We performed western blot analysis utilizing a polyclonal antibody for PACT/RAX to detect the truncated lear-5J protein (Figure 5 A). wt PACT/RAX has a molecular weight of 34 kDa (lanes 1 and 4, upper band), whereas, the lear-5J truncated protein has a predicted molecular weight of 22 kDa (lower band, lanes 2 and 3). Lane 1 shows that in extract derived from *Prkra^lear-5J^*heterozygous mice both wt PACT/RAX (upper band) and lear-5J (lower band) are detectable (lane 1). As expected, extracts prepared from two independent *Prkra^lear-5J^*homozygous mouse brains have no detectable wt PACT/RAX band at the 34 kDa, however, both have a detectable band at 22 kDa (lanes 2-3). Finally, we detect the presence of wt PACT/RAX but not the lear-5J truncated protein in extract derived from brains of wt (BTBR T^+^ Itpr^tf^/J) mice (lane 4). Blots were then probed for β-actin to ensure equal protein loading (lower panel, lanes 1-4). To further validate that the bands observed in figure 5 A are specific to lear-5J and the mRNA corresponding to the *Prkra* gene is present in the brain (and not degraded via NMD), we performed reverse transcriptase PCR (RT-PCR) on total RNA isolated from the brains of these mice (Figure 5 B). Our results indicate that compared to the wt mice (lane 1) the *Prkra* transcripts are present at reduced levels in *Prkra^lear-5J^* mice (lanes 2-3) with the greatest reduction observed in *Prkra^lear-5J^* homozygous mice (lane 3). These results indicate that although NMD may be operative on lear-5J mRNA, it it does not eliminate the transcripts from *Prkra* gene in *Prkra^lear-5J^* brains. This also confirms that the protein bands observed at the expected position in *Prkra^lear-5J^* brain extracts (Figure 5 A) are the truncated mutant lear-5J protein. RT-PCR for the ribosomal protein S15 was used as a positive control to ensure equal amount of cDNA was added to each PCR reaction (lower panel, figure 5 B).

**Figure 5:**
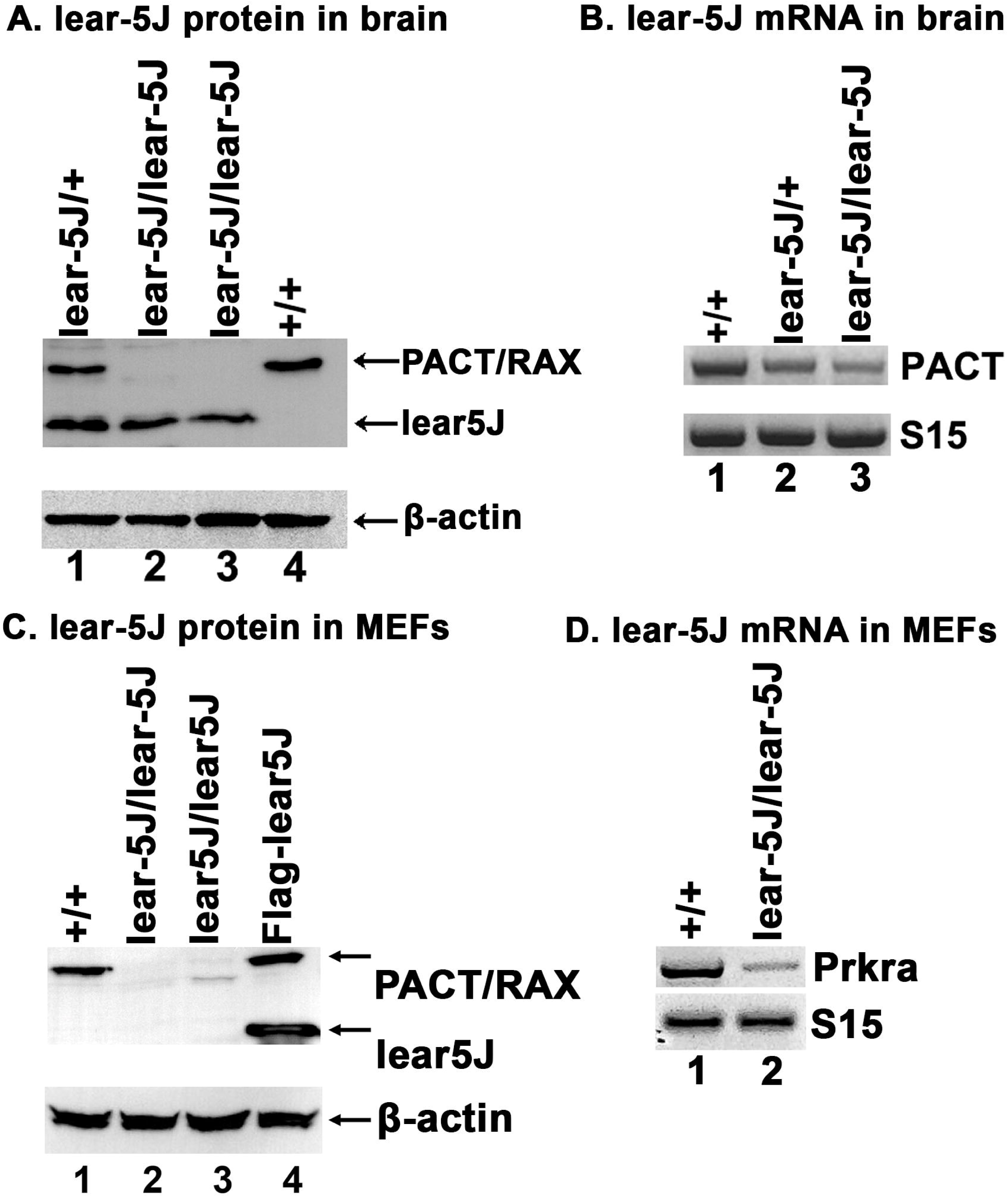
Lear-5J truncated mutant protein is present in mouse brains but not in MEFs. (A) Western blot analysis using brain extracts prepared from wt and *Prkra^lear-5J^*brain samples. Blots were probed for PACT/RAX using a polyclonal antibody and the best of 3 representative blots is shown. The position of the full-length PACT/RAX and the truncated lear-5J protein is indicated by arrows. The blots were probed with β-actin antibody which indicates equal loading. (B) Reverse transcriptase PCR (RT-PCR) using total RNA from the brains of the indicated *Prkra^lear-5J^* genotypes using ribosomal protein S15 as the positive control to ascertain presence of equal amount of total RNA in all samples. (C) Western blot analysis of cell extracts derived from MEFs of the indicated *Prkra^lear-5J^* genotypes and HeLa cells overexpressing Flag-Lear-5J protein. Blots were probed for PACT/RAX using a polyclonal antibody and the best of 3 representative blots is shown. The position of the full-length PACT/RAX and the truncated lear-5J protein is indicated by arrows. (D) RT-PCR using total RNA isolated from wt and homozygous *Prkra^lear-5J^* MEFs. Ribosomal protein S15 was used as a positive control to ascertain equal amount of total RNA was analyzed in each sample.

We next evaluated whether the truncated lear-5J protein was also present in *Prkra^lear-5J^* MEFs. Therefore, we performed western blot and RT-PCR analysis on MEFs derived from wt (BTBR T^+^ Itpr^tf^/J) and lear-5J mice (Figure 5 C and D). HeLa cells over-expressing flag-lear-5J protein were used as a positive control and as a size marker for the lear-5J protein. Our results indicate that wt PACT/RAX is abundant in wt MEFs (lane 1), however, no lear-5J mutant protein is detectable in *Prkra^lear-5J^* MEFs (lanes 2-3). To ensure equal protein loading we then probed for β-actin as our loading control (lower panel, lanes 1-4). Finally, we wanted to determine if the observed absence of lear-5J protein in MEFs is due to the absence of *Prkra* mRNA, as it may be degraded completely via NMD in MEFs. We assessed this using RT-PCR as described before. Our results (Figure 5 D) show a dramatic reduction in detectable *Prkra* mRNA in MEFs homozygous for the *Prkra^lear-5J^* mutation (lane 2) compared to the wt (BTBR T^+^ Itpr^tf^/J) control (lane 1). Ribosomal protein S15 was used as the positive control to ensure equal quantities of cDNA were used for each reaction. These results indicate that NMD may be more efficient in MEFs than in brain. In addition, these results indicate that the truncated mutant lear-5J protein may have reduced stability in MEFs as no lear-5J protein is detected in *Prkra^lear-5J^* MEFs.

### PACT/RAX is expressed in the mouse cerebellum

Previous studies have demonstrated that PACT/RAX contributes to craniofacial development (Dickerman et al., 2011; Palmer et al., 2016a) and for migration of cerebellar granule neurons (Yong et al., 2015). As the *Prkra^lear-5J^*mouse exhibits dystonia, we next evaluated the presence of PACT/RAX in the cerebellum of a developing wt (BTBR T^+^ Itpr^tf^/J) and *Prkra^lear-5J^*mice. The cerebellum is known for motor coordination and proprioception and a growing number of studies have indicated a pathophysiological role of the cerebellum in dystonia (Bologna and Berardelli, 2018; Jinnah and Hess, 2018; Kaji et al., 2018; Gill and Sillitoe, 2019; Morigaki et al., 2021; Gill et al., 2023). Our results show that PACT/RAX is abundantly expressed in the cerebellum (figure 6 A and B). Notably, we observe the highest concentration of PACT/RAX in the Purkinje neuron layer (Figure 6 A, brown staining, and Figure 6 B red fluorescence). Considering this observation, we further evaluated the expression of PACT/RAX in the Purkinje neurons by performing double immunostaining for the Purkinje neuron specific marker, calbindin (green), and PACT/RAX (red), with nuclear stain DAPI (blue) (figure 6 B). The co-localization of red and green fluorescence and presence of yellow fluorescence in the Purkinje neurons confirms that PACT/RAX protein is expressed at high levels in these neurons.

**Figure 6.**
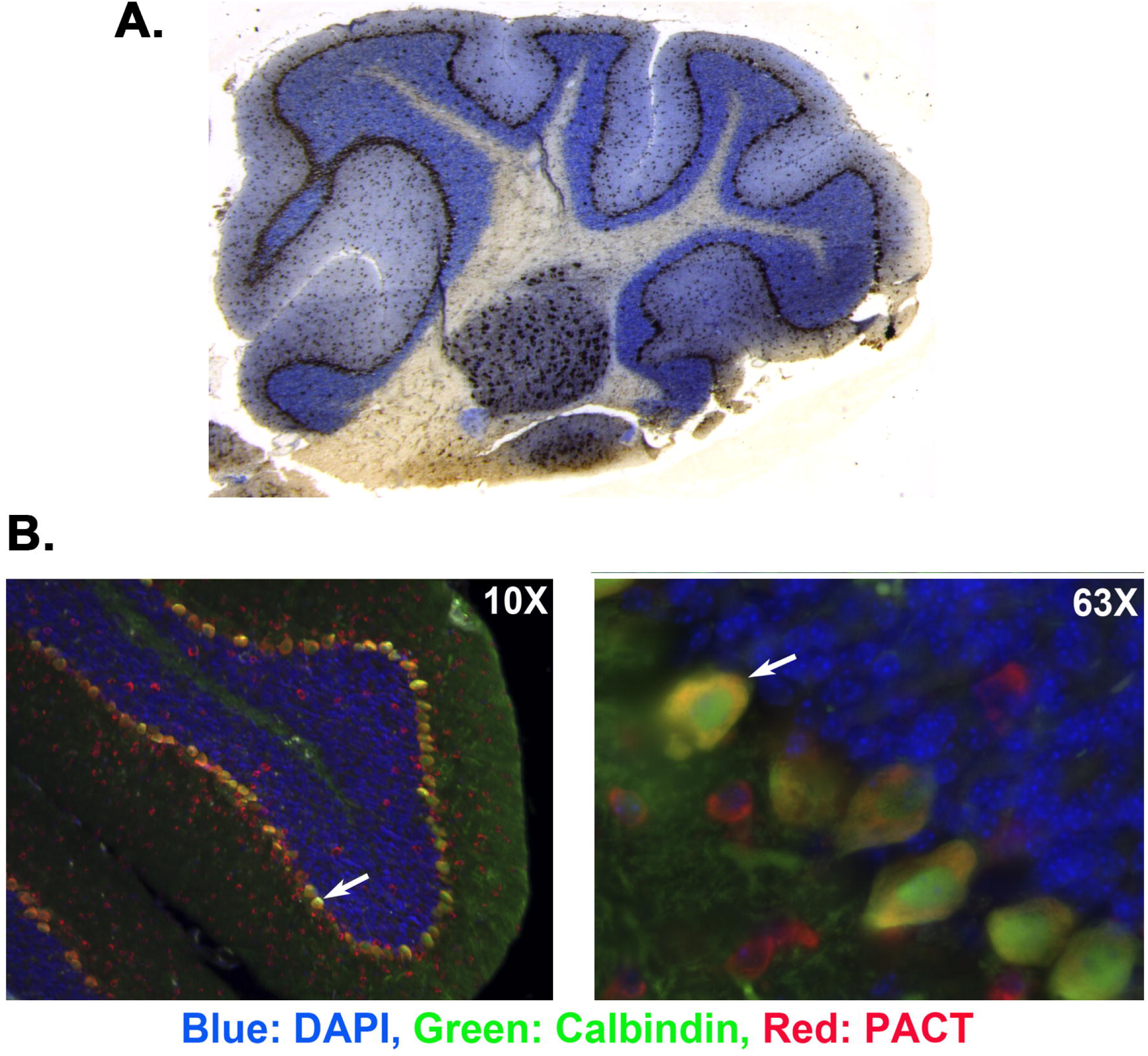
PACT/RAX/RAX protein is abundantly expressed in mouse cerebellum and especially in Purkinje neurons. (A) Immunohistochemistry on day 28 sagittal section of wt C57BL/6 cerebellum using anti-PACT/RAX/RAX antibody. Brown staining indicates presence of PACT/RAX protein. (B) Immunohistochemistry of tissue described in B showing co-localization of PACT/RAX staining with a Purkinje neuron-specific marker calbindin. PACT/RAX (red), DAPI (blue), and calbindin (green).

### *Prkra^lear-5J^* mice show defects in cerebellar development and deficiencies in Purkinje neuron arborization

The cerebellum of mammals is divided into 10 distinct folia (Sillitoe and Joyner, 2007) and after establishing that PACT/RAX is expressed in the mouse cerebellum, we assessed if the *Prkra^lear-5J^* frameshift mutation has any effect on cerebellar folia. To address this, we compared mid-sagittal sections of wt (BTBR T^+^ Itpr^tf^/J) and *Prkra^lear-5J^* cerebellum derived from 28-day old mice (2 of each genotype) using Hematoxylin and Eosin (H&E) (figure 7 A and B). In both wt (BTBR T^+^ Itpr^tf^/J) and *Prkra^lear-5J^* mice we observed the characteristic 10 folia within the cerebellum, but the complexity of the folia was reduced in the *Prkra^lear-5J^*mice. The *Prkra^lear-5J^*folia demonstrate an elongated and less branched pattern relative to that of the BTBR T^+^ Itpr^tf^/J mice. The most distinguishing factor when comparing these foliations is within folium IX. In mice, this folium is subdivided into three lobules (IXa, IXb, IXc)(Sudarov and Joyner, 2007). While we can observe these distinct lobules in the cerebellar tissue derived from wt mice, we do not detect such lobules in the *Prkra^lear-5J^* cerebellum. These initial observations indicate that the cerebellar development of *Prkra^lear-5J^*mice is affected, while noting that both *Prkra^lear-5J^* brains that we evaluated were drastically smaller than the control mice. Due to the abundance of PACT/RAX in the Purkinje cell layer seen in figure 6, we next determined if there were any significant differences in Purkinje neurons between wt (BTBR T^+^ Itpr^tf^/J) and *Prkra^lear-5J^* mice. We performed immunostaining on sagittal sections of the cerebellar tissue using calbindin antibody (green) to specifically mark Purkinje neurons and DAPI (blue) as the nuclear stain (figure 7). Overall organization of Purkinje neurons were similar in both cerebellum sections (Fig. 7 C and E). In wt (BTBR T^+^ Itpr^tf^/J) mice, Purkinje neurons demonstrate the characteristic well-branched arborization from the cell body (figure 7 D). However, in the *Prkra^lear-5J^*cerebellum, we identified a dramatic reduction in the dendritic branching of Purkinje neurons (figure 7 F) compared to the controls.

**Figure 7.**
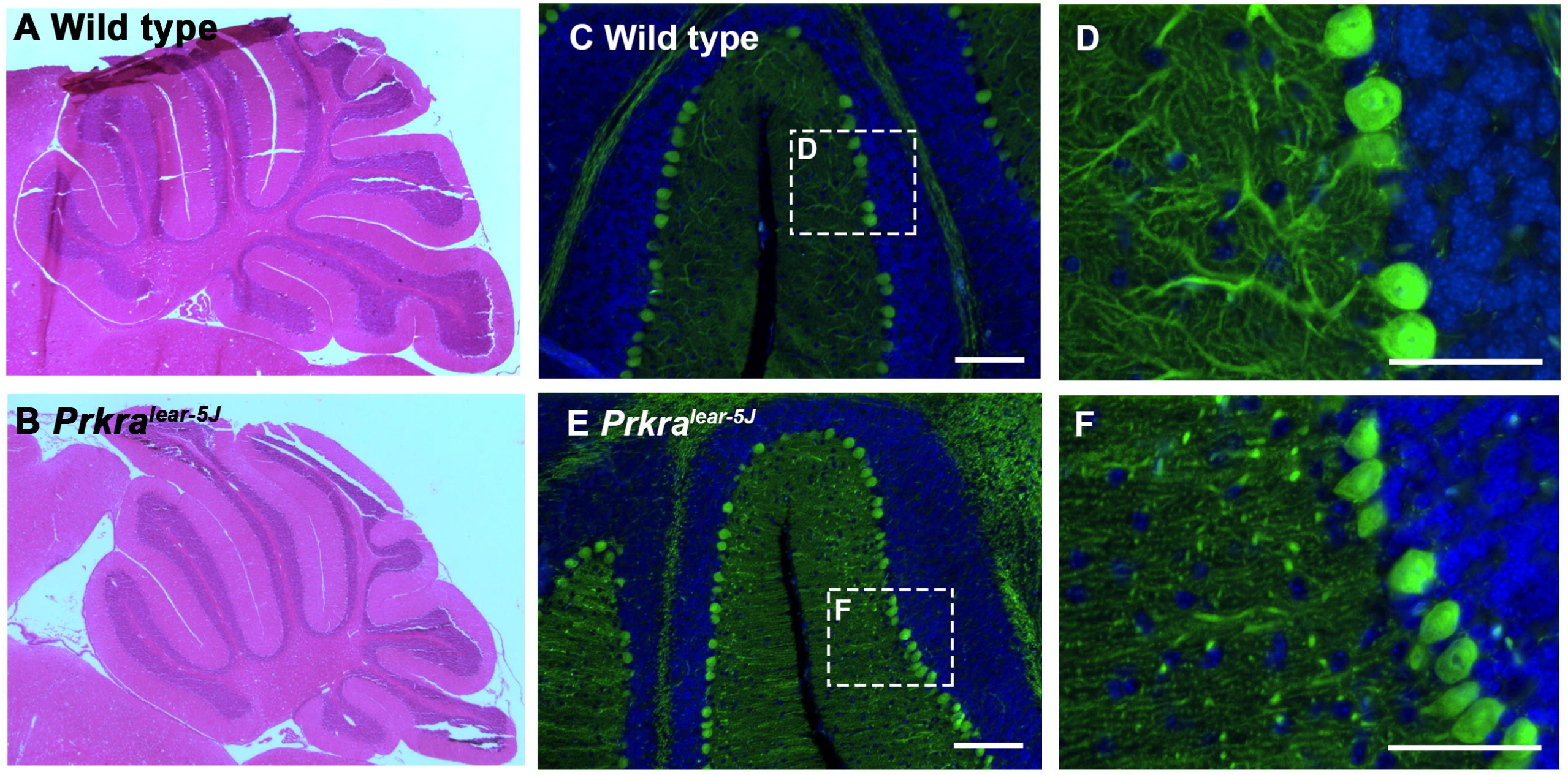
The *Prkra^lear-5J^* mutation affects cerebellar development and reduces arborization in Purkinje neurons. (A-B) Hematoxylin and eosin staining on day 28 sagittal sections of wt (BTBR T^+^ Itpr^tf^/J) and *Prkra^lear-5J^* cerebellum. Fully developed mouse cerebellum has ten lobules denoted here as I-X. (C-F) Immunohistochemistry of Day 28 sagittal sections of mouse cerebellar tissue stained with Purkinje neuron marker, calbindin (green), and nuclear stain DAPI (blue). Dashed boxes (left panel) indicate areas of magnification (right panel).

### *Prkra^lear-5J^* cerebellum shows a significant reduction in eIF2α phosphorylation

A perturbation of the basal eIF2α phosphorylation levels as well as dysregulation of stress-induced phosphorylation has been previously linked to DYT-PRKRA (Vaughn et al., 2015; Burnett et al., 2019; Burnett et al., 2020) and also to other types of dystonia (Rittiner et al., 2016; Beauvais et al., 2018; Zakirova et al., 2018). As PACT/RAX mediated regulation of PKR kinase activity impacts eIF2α phosphorylation, we next determined if the *Prkra^lear-5J^* mutation resulted in any changes in eIF2α phosphorylation in the mouse cerebellum. To answer this question, we performed immunostaining on mid-sagittal sections of mouse cerebellum probing with antibodies that specifically detect phosphorylated eIF2α, and CreP the constitutively expressed phosphatase that dephosphorylates eIF2α. Our results indicate that mice homozygous for the *Prkra^lear-5J^*mutation show a dramatic reduction in levels of phosphorylated eIF2α as compared to wt (BTBR T^+^ Itpr^tf^/J) controls (figure 8 A and B). Interestingly, our results indicate that the *Prkra^lear-5J^* mice have elevated CreP levels as compared to the wt controls (figure 8 C and D) and this may contribute to the presence of low levels of phosphorylated eIF2α in addition to inhibition of PKR by the lear-5J protein. Furthermore, the western blot analysis shown in Fig. 8 B confirms that cerebellar extracts from the *Prkra^lear-5J^* homozygous mice (lanes 4-7) have significantly less eIF2α phosphorylation compared to the extracts from the wt (BTBR T^+^ Itpr^tf^/J, lanes 1 and 2). Additionally, the levels of CreP are higher in cerebellar extracts from *Prkra^lear-5J^* homozygous mice (lanes 4-7) compared to the wt (lanes 1and 2). Interestingly, the *Prkra^lear-5J^*heterozygous cerebellar extract (lane3) shows eIF2α phosphorylation levels higher than both the wt (lanes 1 and 2) and *Prkra^lear-5J^* homozygotes (lanes 4-7). The reasons for this are currently unclear and will require additional analysis. Taken together with the results in Figs. 3 and 8 A, this indicates that the lower levels of eIF2α phosphorylation in *Prkra^lear-5J^* cerebellum could result due to inhibition of PKR by *Prkra^lear-5J^* protein as well as higher expression of CreP.

**Figure 8.**
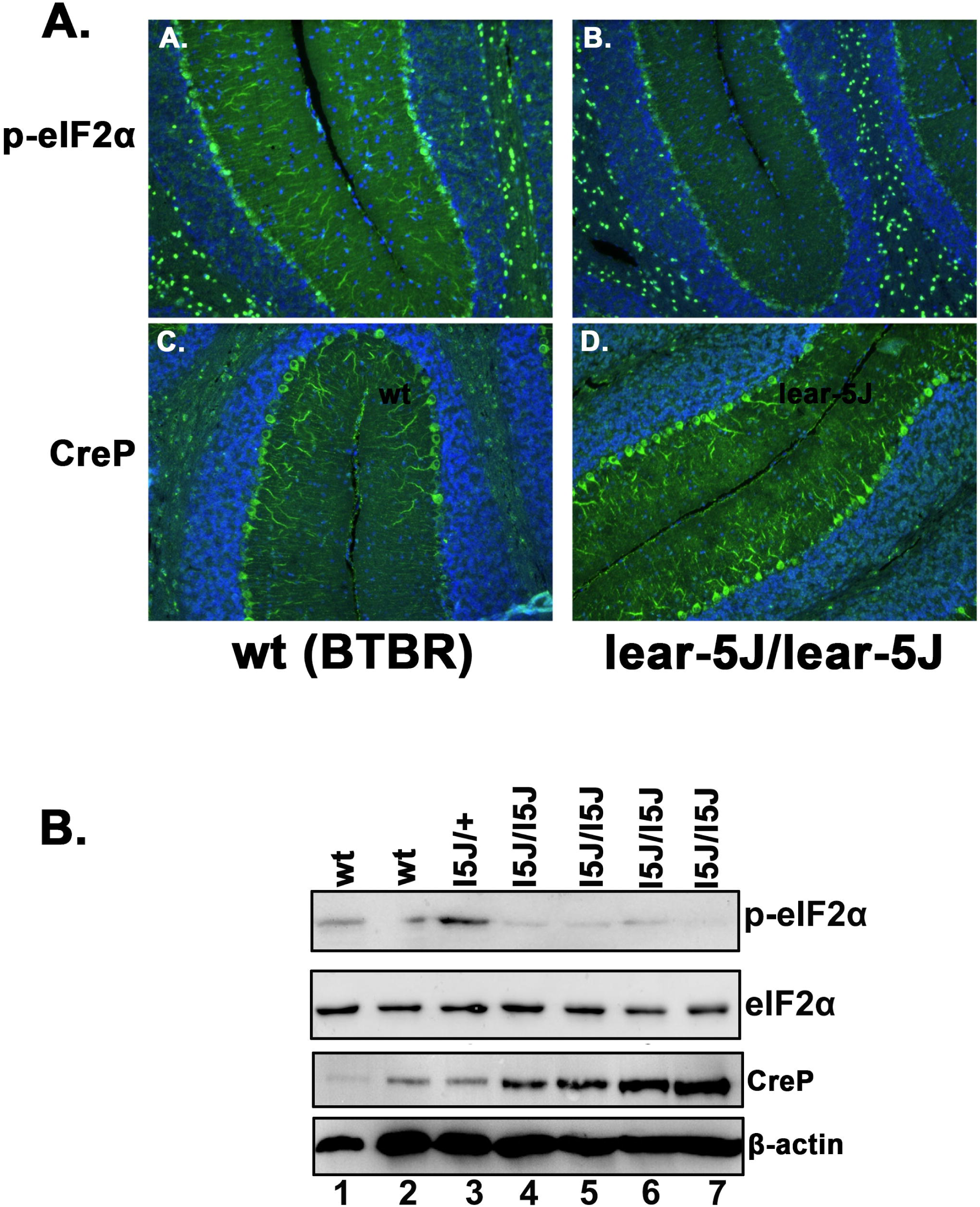
*Prkra^lear-5J^*cerebellum exhibits dysregulation of eIF2α phosphorylation. A. Immunohistochemistry of Day 28 sagittal sections of wt (BTBR T^+^ Itpr^tf^/J) (left) and *Prkra^lear-5J^*(right) mouse cerebellum stained with the nuclear marker DAPI (blue) and protein of interest (green). A and B: DAPI (blue) and phosphorylated eIF2α (green), C and D: DAPI (blue) and CreP (green). B. Western blot analysis of cerebellar extracts. Extracts prepared from 2 wt (BTBR T^+^ Itpr^tf^/J) (lanes 1-2), 1 *Prkra^lear-5J/+-^* heterozygous (lane3) and 4 *Prkra^lear-5J/lear5-J^* homozygous cerebellums (lanes 4-7) were analyzed with antibodies specific for p-eIF2α, total eIF2α, CreP, and β-actin. B. Western blot analysis of wt (BTBR T^+^ Itpr^tf^/J) and *Prkra^lear-5J^* cerebellar extracts. Cerebellar extracts prepared from two wt (BTBR T^+^ Itpr^tf^/J), one heterozygous *Prkra^lear-5J/+^*, and four homozygous *Prkra^lear-5J/lear-5J^*mice were analyzed by western blot analysis. Blots were probed for p-eIF2α, total eIF2α, and CreP phosphatase using specific antibodies and the best of 3 representative blots is shown. The blots were probed with β-actin antibody to ensure equal loading.

## Discussion

PACT/RAX serves as a negative regulator of general protein synthesis under conditions of cellular stress by triggering eIF2α phosphorylation via PKR activation (Ito et al., 1999; Patel et al., 2000; Bennett et al., 2004; Bennett et al., 2006; Singh et al., 2009; Chukwurah and Patel, 2018). Our previous research has established that several DYT-PRKRA mutations lead to enhanced PKR activation and eIF2α phosphorylation in response to ER stress. The recessively inherited P222L mutation leads to a delayed but heightened and persistent PKR activation and eIF2α phosphorylation to increase cell susceptibility to ER stress (Vaughn et al., 2015). A dominantly inherited frameshift mutation causes insoluble protein aggregates, PKR activation, and significant apoptosis even in the absence of stress (Burnett et al., 2019). Additionally, 3 other recessive and 2 dominant mutants also exhibit enhanced PKR interaction and activation causing increased eIF2α phosphorylation and sensitivity to ER stress in DYT-PRKRA patient cells (Burnett et al., 2019). In addition to our studies on DYT-PRKRA, maladaptive eIF2α signaling is also seen in DYT-TOR1A (DYT1), DYT-THAP1 (DYT6), and sporadic cervical dystonia (Rittiner et al., 2016; Beauvais et al., 2018; Zakirova et al., 2018). Thus, dysregulated eIF2α phosphorylation has emerged as a convergent theme in dystonia.

In the present study, our objective was to determine if the murine *Prkra^lear-5J^* mutation results in dysregulated eIF2α signaling contributing to the dystonia-like symptoms. A characterization of the biochemical properties on the truncated lear-5J protein revealed that it has reduced ability to bind dsRNA while retaining its ability to interact with PKR. This was expected, as we have previously characterized that the high affinity binding to dsRNA requires two intact dsRBMs (Huang et al., 2002) and the lear-5J protein has lost the carboxy terminal part of dsRBM2 critical for binding to dsRNA. Based on our previous research, as the interaction of PACT/RAX with PKR is primarily mediated via the dsRBM1 (Chukwurah et al., 2018), the lear-5J protein retains its ability to interact with PKR. For PACT/RAX to activate PKR, the intact M3 motif is essential and we and others previously characterized that a deletion of M3 renders PACT/RAX unable to activate PKR despite its ability to interact with PKR (Peters et al., 2001; Huang et al., 2002). Thus, it is not surprising that lear-5J protein has lost its ability to activate PKR although it can interact with PKR. Additionally, lear-5J protein inhibits PKR in the presence of both activators, dsRNA or PACT/RAX, which is likely due to a direct binding of lear-5J protein to PKR, thus preventing its interaction with dsRNA or PACT/RAX. Alternately, lear-5J protein could also directly bind to dsRNA and wt PACT/RAX protein to prevent their interaction with PKR resulting in the observed inhibition of PKR kinase activity.

The presence of lear-5J protein in the brain but not in MEFs indicates that the truncated mutant protein may be degraded efficiently in MEFs but not in the brain. The lear-5J mRNA could be detected both in brain and MEFs indicating that it is not degraded efficiently by NMD in both tissues. The efficiency of NMD is dependent on many factors depending on the location of the nonsense mutation, cell type, and the mutated gene (Miller and Pearce, 2014). Thus, it is not surprising to detect the mRNA encoding lear-5J protein at variable levels in the brain and MEFs. It is worth noting that we have previously characterized the PACT/RAX null MEFs to be significantly resistant to apoptosis in response to ER stressor tunicamycin (Singh et al., 2009). Resistance of *Prkra^lear-5J^*MEFs to ER stress observed in the current study thus agrees with previous studies that the absence of PACT/RAX confers resistance to ER stress-induced apoptosis.

Truncated lear-5J protein being present in the brain may also be one possible reason for the dystonia phenotype observed in the *Prkra^lear-5J^*homozygous mice but not in *Prkra*^-/-^ mice. Rowe *et. al* previously described a *Prkra^-/-^* mouse that was generated by targeting a 3’ exon does not demonstrate any dystonia-like symptoms; however, it shares many overlapping developmental phenotypes with the *Prkra^lear-5J^* mice (Rowe et al., 2006a; Dickerman et al., 2015; Palmer et al., 2016a). Both *Prkra* ^-/-^ and *Prkra^lear-5J^*mice exhibit reduced body size, craniofacial abnormalities, under-developed or small external ears, and hearing loss (Rowe et al., 2006a; Dickerman et al., 2015). Another contributing factor to the lack of a dystonia phenotype in *Prkra*^-/-^ mice can be that it was produced in a different genetic background. The *Prkra* ^-/-^ mice are described on C57BL/6 background, whereas, the *Prkra^lear-5J^* mice are on the BTBR T^+^ Itpr3^tf^/J background (Rowe et al., 2006b; Palmer et al., 2016b). The BTBR T^+^ Itpr3^tf^/J genetic background contains a mutation in the Itpr3 gene which codes for the inositol 1,4,5-triphosphate receptor type 3. This receptor plays a critical role in regulating intracellular calcium levels (Mangla et al., 2020) and as calcium signaling is directly linked to ER homeostasis and eIF2α signaling, it is reasonable to speculate that the combination of calcium signaling dysregulation and *Prkra^lear-5J^* mutation may result in the enhanced dystonia phenotype, especially since we have previously linked PACT/RAX mediated PKR regulation to play a central role in cellular fate in response to ER stress (Singh et al., 2009; Vaughn et al., 2015; Burnett et al., 2019). Previous reports demonstrate that the BTBR mice exhibit abnormal cerebellar development leading to motor dysfunction (Xiao et al., 2020). This study noted an increased proliferation of cerebellar granule neurons with enhanced cerebellar foliation, and hypertrophy of Purkinje cells with increased dendritic spines in BTBR mice. The cerebellar defects observed in *Prkra^lear-5J^* mice differ markedly from those observed in the BTBR strain, suggesting that the truncated PACT/RAX protein has a specific effect.

Lack of PACT/RAX function was also investigated by Bennett *et al*. in flies and mice (Bennett et al., 2008). Additionally, a mouse missense mutation S130P in the second dsRBM of PACT/RAX exhibited defects in ear development, growth, craniofacial development, and ovarian structure but no movement disorder phenotype in mice with a C57BL/6J and C3HeB/FeJ mixed background (Dickerman et al., 2011). These results indicate that the genetic background can significantly alter the neuromuscular effects of PACT/RAX mutations. In future studies, it will be beneficial to breed the *Prkra^lear-5J^* mutation, as well as other DYT-PRKRA mutations, in a C57BL/6 genetic background which could clarify if the resulting dystonia phenotype arises due to the truncated PACT/RAX protein alone or a combination of *Prkra^lear-5J^* mutation and the BTBR T^+^ Itpr3^tf^/J genetic background. As PACT/RAX is an evolutionarily conserved protein, its function has been studied in *Drosophila*. Flies carrying an inactivating transposon insertion in *Drosophila PACT/RAX* (loqs/R3D1) caused highly abnormal commissural axon structure of the CNS and a severe defect in neuromuscular coordination (Bennett et al., 2008), thus indicating that PACT/RAX may regulate motor coordination via mechanisms other than regulation of PKR as there is no PKR homolog in flies.

The cerebellum is a region of the brain that is indispensable for motor control (Timmann et al., 2010; Welniarz et al., 2021). When comparing the overall cerebellar developmental pattern of the *Prkra^lear-5J^* mice to that of wt (BTBR T^+^ Itpr3^tf^/J) mice, an overall reduction in the complexity of the foliation and a severe deficiency in the arborization of the Purkinje neurons is noted. Purkinje neurons facilitate communication between various cerebellar regions to control the final output in regulation of dexterous limb movements through their extensive dendritic arborization (Thanawalla et al., 2020; Gaffield et al., 2022). Interestingly, previous studies have specifically identified that the dysregulation of either sodium or potassium pumps in Purkinje neurons leads to the rapid onset of dystonia and parkinsonism or ataxia (Calderon et al., 2011; Tara et al., 2018). Our results suggest that the Purkinje neurons in *Prkra^lear-5J^* mice may have compromised functionality because of the severe deficit of dendritic arborizations, and that could contribute to the underlying dystonia phenotypes. Interestingly, perturbations in eIF2α phosphorylation has been linked to a dysregulation of dendritic arborization in flies (Tsuyama et al., 2017). Previously, it has also been reported that PACT/RAX is essential for the migration of cerebellar granule neurons in the developing mouse cerebellum (Yong et al., 2015), thus indicating that PACT/RAX may regulate cerebellar development at multiple levels.

The *Prkra^lear-5J^*mice have a reduction in the basal eIF2α phosphorylation levels and elevated levels of CreP phosphatase (Hicks et al., 2023). The inhibition of PKR activity by lear-5J truncated mutant protein and increased CreP levels could both contribute to the observed reduction in basal eIF2α phosphorylation. This contrasts with the increased phosphorylation of eIF2α phosphorylation, heightened PKR kinase activity and enhanced sensitivity to ER stress in DYT-PRKRA patient cells (Vaughn et al., 2015; Burnett et al., 2020). Additionally, similar increases in PKR activity and eIF2α phosphorylation was reported in DYT-EIF2AK2 (DYT33) patients carrying PKR missense variants with early onset generalized dystonia (Kuipers et al., 2021). However, preventing or reducing eIF2α phosphorylation is also detrimental for coordinated muscle movement. The most compelling and recent evidence comes from studies on DYT-TOR1A (DYT1), where compounds that boosted the ISR showed corrective activity in cell and animal models (Caffall et al., 2021). Thus, boosting eIF2α phosphorylation could work in alleviating dystonia symptoms in cases where ISR is attenuated such as DYT1 and sporadic cervical dystonia. It is also important to note that in case of DYT-EIF2AK2 (DYT33), PKR inactivating mutations were also reported in some patients (Mao et al., 2020), thereby suggesting that a reduction in PKR activity and consequently reduced eIF2α phosphorylation may also lead to dystonia pathophysiology. Furthermore, a selective deletion of PERK, one of the eIF2α kinases, in mouse midbrain dopaminergic neurons resulted in multiple cognitive and age-dependent motor phenotypes which could also be observed by expression of a phospho-mutant eIF2α (Longo et al., 2021). It is likely that precise regulation of the extent and duration of eIF2α phosphorylation is essential for optimal neuronal regulation of motor control and either a reduction or elevation of the ISR response may lead to lack of motor coordination. Recently, it was reported that the cholinergic neurons constitutively engage the ISR for dopamine modulation and skill learning (Helseth et al., 2021). Although phosphorylation of eIF2α has classically been viewed as a stress response, eIF2α phosphorylation mediated regulation of protein synthesis is utilized by neurons for mechanisms besides stress response that include behavior, memory consolidation, neuronal development, and motor control (Bellato and Hajj, 2016). Future research using targeted mutations in specific neuronal subtypes can test the exact contribution of ISR and specifically eIF2α phosphorylation for neuronal control of muscle movement.

In conclusion, we identify that reduced eIF2α phosphorylation resulting from a frameshift mutation in murine PRKRA gene affects cerebellar development. This study combined with our earlier studies with DYT-PRKRA patient cells provide direct evidence for a key role of a dysfunctional eIF2α pathway in the pathogenesis of dystonia. The *Prkra^lear-5J^*mice also can be a suitable mouse model for screening potential therapeutic options to alleviate the dystonia symptoms arising from dysregulated eIF2α signaling and should be tested for the suitability for such studies.

## Methods

### Cell Lines and Antibodies

Both HeLaM and Mouse Embryonic Fibroblasts (MEFs) cell lines were cultured in Dulbecco’s Modified Eagle’s Medium (DMEM) containing 10% Fetal Bovine Serum and penicillin/streptomycin. All transfections were carried out using Effectene transfection reagent (Qiagen) per manufacturer protocol. Antibodies used were as follows: PKR: anti-PKR (human) monoclonal (71/10, R&D Systems), p-eIF2α: anti-phospho-eIF2α (Ser-51) polyclonal (CST, #9721), PACT: anti-PACT polyclonal (ProteinTech, 10771-1-AP), ATF4: Anti-ATF4 monoclonal (CST, #11815), FLAG-HRP: anti-FLAG monoclonal M2-HRP (Sigma A8592), β-Actin: Anti-β-Actin-Peroxidase monoclonal (Sigma-Aldrich, A3854), CreP: PPP1R15B rabbit polyclonal (ProteinTech, 14634-1-AP).

### *Prkra^lear5-J^* mice and tissues

Generation and maintenance of *Prkra^lear5-J^* mice was as described before. Brain tissues were collected from *Prkra^lear5-J^* and BTBR T^+^ Itpr3^tf^/J. The MEFs were generated from E14.5 embryos from *Prkra^lear5-J^* intercrosses.

### Generation of lear-5J Frameshift Mutation

We generated the lear-5J frameshift mutation using site specific mutagenesis through PCR amplification such that antisense primer contained a single adenine insertion relative to the *Prkra* template DNA. The following site-specific mutagenic primers were used: lear-5J Sense: GCCTCGAGCACATATGTCCCATAGCAGGCATCG lear-5J Antisense: GCGGATCCGAAATATTACTAAACTTGGCGAGAAATTTCTCAGCAGCATTTCTC TTGGCTTGTTTTTTTTGATGCCCCCTTTC. The subsequent PCR product was then ligated into pGEMT-Easy vector (Promega) and the presence of the mutation was validated through DNA sequencing services provided by Eton Biosciences. After sequence validation we generated amino terminal flag-tagged lear-5J constructs in pBSIIKS+ using NdeI and BamHI restriction sites. We next subcloned the Flag-lear-5J ORF into the mammalian expression construct, pCDNA3.1-, using XbaI-BamHI restriction sites.

### Co-Immunoprecipitation Assays

HeLa M cells were seeded at 20% confluency in 6-well dishes 24-hours prior to transfecting 500 ng of either flag-wt PACT or flag-lear-5J constructs using Effectene transfection reagent (Qiagen). Cells were harvested 24-hours post transfection and harvested in IP buffer (20 mM Tris-HCl pH 7.5, 150 mM NaCl, 1 mM EDTA, 1% Triton X-100, 20% Glycerol). Whole cell extract was then immunoprecipitated overnight at 4°C on a rotating wheel using anti-PKR antibody (71/10, R&D Systems) conjugated to protein A sepharose beads (GE Healthcare) in IP buffer. Immunoprecipitates were then washed 3 times in IP buffer followed by resuspension and boiling for 5 minutes in 2x Laemmli buffer (150 mM Tris–HCl pH 6.8, 5% SDS, 5% β-mercaptoethanol, 20% glycerol). Samples were then resolved on 12% SDS-PAGE denaturing gels and transferred onto PVDF membranes. Blots were initially probed for flag-PACT/lear-5J constructs (coIP) using anti-flag antibody (described above) followed by incubation in anti-PKR antibody (described above) to determine equal amounts of PKR were immunoprecipitated. Input blots of whole cell extract without immunoprecipitation are shown to indicate equal amounts of protein in each sample.

### dsRNA-binding assay

Both wt PACT and lear-5J constructs in pCDNA3.1-were *in vitro* translated using the TNT-T7-coupled rabbit reticulocyte system from Promega while incorporating an ^35^S-Methionine radiolabel and the dsRNA binding ability was measured using poly(I:C) conjugated agarose beads. We diluted 4 μl of *in vitro* translation in 25 μl of binding buffer (20 mM Tris–HCl, pH 7.5, 0.3 M NaCl, 5 mM MgCl_2_, 1 mM DTT, 0.1 mM PMSF, 0.5% NP-40, 10% glycerol) and incubated in 25 μl of poly(I:C)-agarose beads and incubated at 30°C for 30-min. We then washed the beads 4 times with 500 μl of binding buffer and bound proteins were analyzed *via* SDS-PAGE gel electrophoresis and autoradiography. The competition assay was performed incubating either soluble single-stranded RNA, poly(C), or dsRNA, poly(I:C), with the poly(I:C)-agarose beads before the adding the *in vitro* translated proteins. To ensure the presence of PACT was due to the dsRNA binding capacity we assayed *in vitro* translated ^35^S-Methionine labeled firefly luciferase which has no dsRNA binding ability. Bands in bound and total lanes were quantified using Typhoon FLA7000 by analyzing relative band intensities of both T and B lanes. Percentage of PACT bound to beads was calculated and plotted as bar graphs.

### Expression and purification of PACT/lear-5J from E. coli

The ORFs of both wt PACT and lear-5J frameshift mutation were subcloned into pET15b (Novagen) to generate an in-frame fusion protein with a histidine tag. Recombinant proteins were then expressed and purified as previously described (Patel and Sen, 1998).

### PKR Activity Assay

HeLa M cells treated with IFN-β for 24-hours and harvested at 70% confluency. Cells were washed using ice-cold PBS and centrifuged at 600 g for 5-minutes. Cells then resuspended in lysis buffer (20 mM Tris–HCl pH 7.5, 5 mM MgCl_2_, 50 mM KCl, 400 mM NaCl, 2 mM DTT, 1% Triton X-100, 100 U/ml aprotinin, 0.2 mM PMSF, 20% glycerol) and incubated on ice for 5 minutes. Whole cell lysates were then centrifuged at 10,000 g for an additional 5-minutes. PKR was then immunoprecipitated from 100 ug of this lysate using anti-PKR monoclonal antibody (R&D Systems Technology: MAB1980) in a high salt buffer (20 mM Tris–HCl pH 7.5, 50 mM KCl, 400 mM NaCl, 1 mM EDTA, 1 mM DTT, 100 U/ml aprotinin, 0.2 mM PMSF, 20% glycerol, 1% Triton X-100) at 4°C on a rotating wheel for 30-minutes. We then added 10 ul of Protein A-Sepharose beads to each immunoprecipitate followed by an additional 1-hour incubation under the same conditions. Protein A-Sepharose beads were then washed 4 times in high salt buffer followed by an additional two washes in activity buffer (20 mM Tris–HCl pH 7.5, 50 mM KCl, 2 mM MgCl_2_, 2 mM MnCl2, 100 U/ml aprotinin, 0.1 mM PMSF, 5% glycerol). The PKR activity assay using the PKR bound to protein A-Sepharose beads was then conducted by incorporating: 500 ng of purified eIF-2 as the PKR substrate, 0.1 mM ATP, 10 uCi of [γ-^32^P] ATP, and increasing amounts of either recombinant wt PACT or recombinant lear-5J (400 pg – 4 ng) as the PKR activator. Reaction was then incubated at 30°C for 10 min and resolved on a 12% SDS-PAGE gel and analyzed via autoradiography.

### DNA fragmentation analysis

DNA fragmentation analysis was performed as described before (Vaughn et al., 2015). 5 × 10^6^ MEFs established from WT and *Prkra^lear-5J^* mice were treated with 0.5 μg/ml tunicamycin for 48 h followed by DNA fragmentation analysis. To quantify the DNA fragmentation, the fluorescence image was inverted, and the total band intensities in the entire lanes were computed with ImageQuant software on Typhoon FLA 7000 PhosphorImager (GE Healthcare and Life Sciences) and compared with untreated samples as well as between WT and patient cells. The band intensities in WT untreated samples were considered as 1.0, and -fold increases in band intensities with respect to WT untreated samples were calculated and subjected to statistical analysis. A statistical analysis from four different experiments was performed to calculate *p* values to determine significant differences between WT and patient untreated and treated samples.

### Caspase 3/7 activity assays

Both wt and *Prkra^lear-5J^* MEFs were seeded at a concentration of 300,000 cells/ml of medium and treated with a concentration of 5 μg/ml of tunicamycin over a 36-h time course. Samples were collected at indicated time points and mixed with equal parts Promega Caspase-Glo 3/7 reagent (Promega G8090) and incubated for 45 min. Luciferase activity was measured and compared to cell culture medium alone and untreated cells as the negative controls. A statistical analysis from six different experiments was performed to calculate *p* values to determine significant differences between WT and *Prkra^lear-5J^*samples.

### Western Blot Analysis

Protein extracts were prepared from a fraction of the brains of wt mice, mice heterozygous for the *Prkra^lear-5J^* mutation, and mice homozygous for the *Prkra^lear-5J^*mutation using western lysis buffer (20 mM Tris–HCl pH 7.5, 5 mM MgCl_2_, 50 mM KCl, 400 mM NaCl, 2 mM DTT, 1% Triton X-100, 100 U/ml aprotinin, 0.2 mM PMSF, 20% glycerol) containing a 1:100 dilution of protease inhibitor (Sigma) and phosphatase inhibitor (Sigma). Tissue was initially homogenized in the lysis buffer cocktail described above and incubated on ice for 5 minutes followed by centrifugation at 13,000 g for 10 minutes. Concentration of total protein extract was then determined through Bradford assays and appropriate amounts were resolved on SDS-PAGE denaturing gels to detect the proteins of interest. Protein extract derived from MEFs were prepared from wt and *Prkra^lear-5J^* MEFs cultured as described above. Cells were initially seeded at 40% confluency and harvested 24-hours later using the lysis buffer cocktail described above. Concentration of whole cell extracts were determined by Bradford assays. Proteins were detected via chemiluminescence and antibodies indicated above.

### Reverse Transcriptase PCR

RNA was isolated from either: (i) a fraction of mouse brain derived from wt mice, *Prkra^lear-5J^* heterozygous mice, *Prkra^lear-5J^* homozygous mice or (ii) wt and lear-5J homozygous MEFs. In all cases, tissue or cells were incubated on ice for 5-minutes in RNA-Bee (Tel-Test, Inc) and in the presence of CIA (chloroform:isoamylacetate (24:1)). Lysates were then centrifuged at 12,000 g for 15-minutes at 4°C. The fraction of lysate containing RNA was then carefully collected and precipitated overnight in an equal volume of isopropanol at -20°C. Samples were then centrifuged at 12,000 g for 15 minutes to pellet the RNA. Supernatant was removed and RNA pellet was washed 2x in 70% ETOH and centrifuged at 12,000g for 8 minutes at 4°C followed by a 1-hour incubation at room temperature to ensure RNA was devoid of alcohol contamination. Purified RNA pellets were then resuspended in nuclease free water. The cDNA was generated using 1 μg of total RNA, 75 nG of random hexamer, and reverse transcriptase (Thermo Scientific) per manufacturer protocol. Finally, relative mRNA levels of *Prkra* were compared using *Prkra* specific primers and ribosomal protein S15 primers were used to amplify this housekeeping gene to ensure equivalent amounts of cDNA were used in each reaction. The following primers were used: *Prkra* Sense: ATGTCCCATAGCAGGCATCGTGCCG, *Prkra* Antisense: CCTTCCTGGGAAAGGGTATATTCAGG, S15 Sense: TTCCGCAAGTTCACCTACC, S15 Antisense: CGGGCCGGCCATGCTTTACG.

### Histology and Immunostaining

Cerebellums derived from day 28 *Prkra^lear-5J^* homozygous and wt (BTBR T^+^ Itpr^tf^/J) mice were fixed at 4°C overnight in 4% paraformaldehyde in PBS. The fixative was then removed, and cerebellums were then washed in PBS followed by dehydration in absolute ethanol. Following the dehydration step, tissues were then permeabilized with methyl salicylate and embedded in paraffin. We then generated 6 μM sagittal sections from each tissue and evaluated through hematoxylin and eosin (H&E) staining. Immunostaining was carried out as previously described (Youngblood et al., 2018) and antibodies, dilutions, and detection methods are indicated below. In brief, experiments utilizing mouse primary antibodies utilized blocking reagents from the M.O.M. kit (Vector Laboraties) and experiments utilizing primary antibodies from other species used the blocking reagent from Tyramide Signal Amplification (TSA) kit no. 22 dissolved in PBS (Thermo Fisher Scientific) each per manufacturer protocol. In each case, primary antibodies were incubated overnight at 4°C in the appropriate blocking reagent. Samples were then incubated in species specific secondary antibodies in TSA block for 30 minutes at room temperature. Biotin-conjugated secondary antibodies were incubated in streptavidin conjugates (Alexa Fluor 488, Alexa Fluor 594, or HRP) at a 1:100 dilution in TSA block at room temperature for 30 minutes (Thermo Fisher Scientific). Following this incubation, the streptavidin-HRP conjugates were either subjected to SigmaFast 3,3’-diaminobenzidine (DAB) staining (Sigma-Aldrich) or TSA with Alexa Fluor 488 (TSA kit no.22, Thermo Fisher Scientific) for 3 minutes in each case. Fluorescent detection methods were counterstained with 300 nM 4’,6-diamidino-2-phenylindole (DAPI) for 5 minutes prior to mounting in fluorescence mounting media. DAB detection was counterstained with 0.5% methyl green in 0.1 M sodium acetate buffer (pH 4.2) for 5 minutes at room temperature.

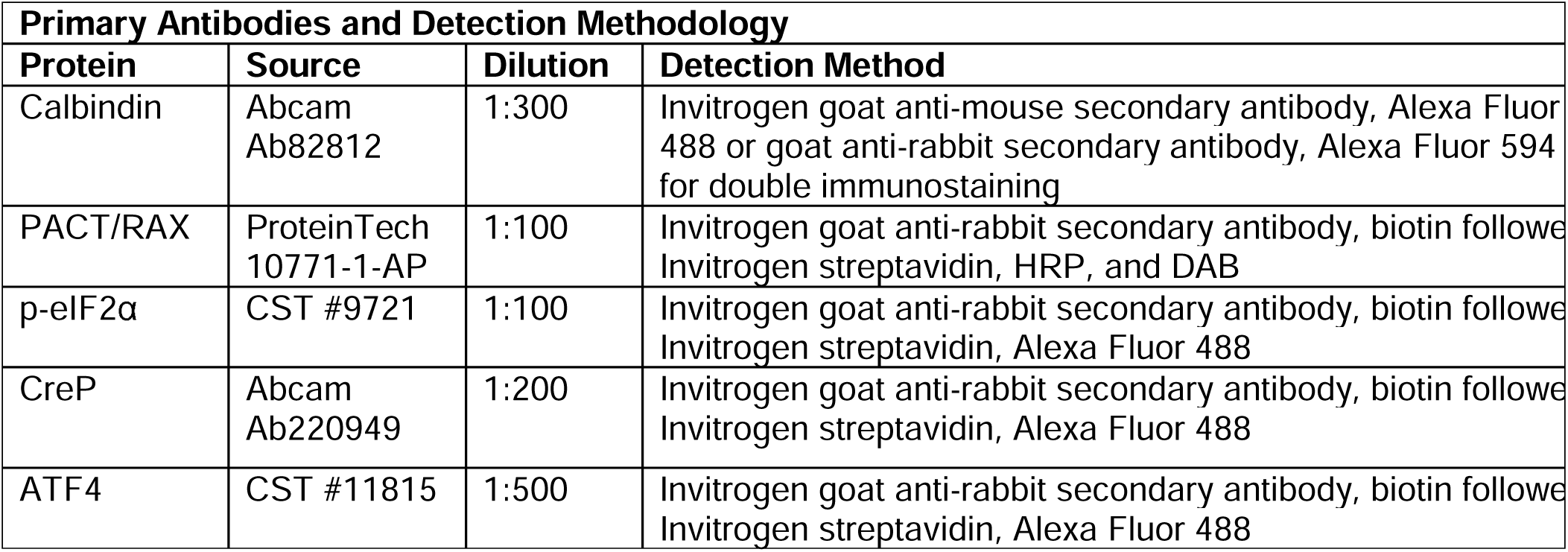

## Notes

Funding Sources: This work was supported by a Department of Defense through the grants W81XWH-18-1-0088, W81XWH-22-1-0526 to RP, a NIHG grant DE020052 to SAM, a SPARC Grant to K F, and an ASPIRE grant to RP from University of South Carolina Vice President’s Office. Opinions, conclusions, and recommendations are those of the author and are not necessarily endorsed by the Department of Defense.

### Competing Interest Statement

The authors have declared no competing interest.

